# Orexin A excites the rat olivary pretectal nucleus via OX_2_ receptor in a daily manner

**DOI:** 10.1101/2021.05.11.443625

**Authors:** Lukasz Chrobok, Anna Alwani, Kamil Pradel, Jasmin Daniela Klich, Marian Henryk Lewandowski

## Abstract

Pronounced environmental changes between the day and night forced living organisms to evolve specialised mechanisms organising their daily physiology, named circadian clocks. Currently, it has become clear that the master clock in the suprachiasmatic nuclei of the hypothalamus is not an exclusive brain site to generate daily rhythms. Indeed, several brain areas, including the subcortical visual system have been recently shown to change their neuronal activity across the daily cycle. Here we focus our investigation on the olivary pretectal nucleus (OPN) – a retinorecipient structure primarily involved in the pupillary light reflex. Using the multi-electrode array technology *ex vivo* we provide evidence for OPN neurons to elevate their firing during the behaviourally quiescent light phase. Additionally, we report the robust sensitivity to orexin A via the identified OX_2_ receptor in this pretectal centre, with higher responsiveness noted during the night. Interestingly, we likewise report a daily variation in the response to PAC1 receptor activation, with implications for the convergence of orexinergic and visual input on the same OPN neurons. Altogether, our report is first to suggest a daily modulation of the OPN activity via intrinsic and extrinsic mechanisms, organising its temporal physiology.

**GRAPHICAL ABSTRACT:** 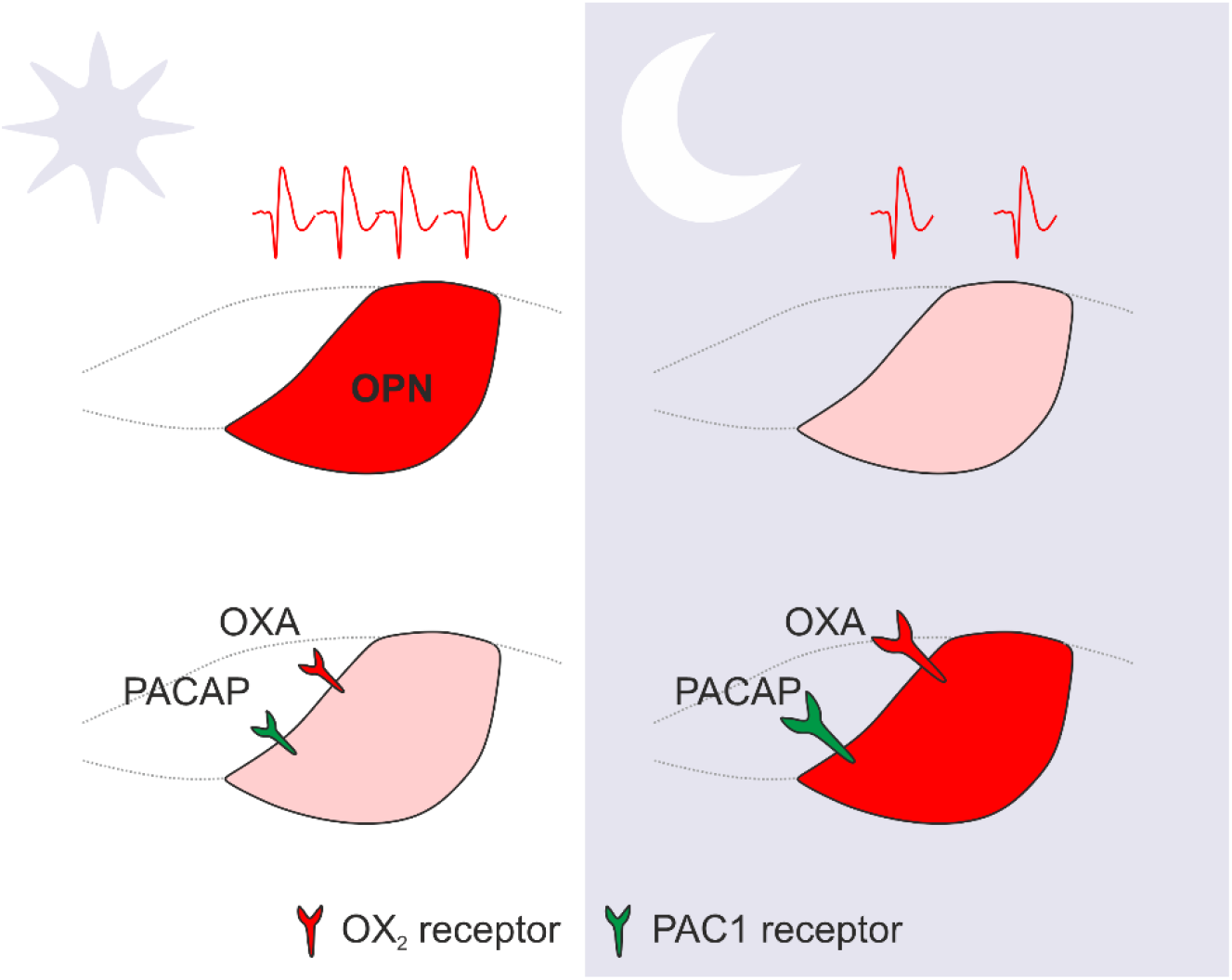

**HIGHLIGHTS:**

- Neurons in the olivary pretectal nucleus (OPN) increase their firing during the day
- Orexin A robustly excites the OPN via the OX_2_ receptor
- Orexin A and the activation of PAC1 receptor are more effective during the night

## 1. INTRODUCTION

Biological clocks evolved to adapt animals to cyclic changes in the environment, with the cyclic changes in the ambient light levels being the most robust and prominent rhythm of the outside world. In mammals, the master clock is localised in the suprachiasmatic nucleus (SCN) of the hypothalamus (Hastings et al., 2019, 2018; Takahashi, 2017). However, recent advances show several other brain areas and peripheral tissues to exhibit autonomous or semi-autonomous circadian rhythms (Begemann et al., 2020; Guilding and Piggins, 2007; Paul et al., 2020). Moreover, the orexinergic system of the lateral hypothalamus has been schematised to act as the hands of the clock; due to its reciprocal connection with the SCN orexins deliver arousal for a plethora of brain sites in a circadian manner (Azeez et al., 2018; Belle et al., 2014; Marston et al., 2008).

The olivary pretectal nucleus (OPN) is a small retinorecipient area which constitutes a key element of the pupillary light reflex (Young and Lund, 1998). Similarly to the master clock, the majority of retinal fibres that reach OPN belong to intrinsically photosensitive retinal ganglion cells (ipRGCs) (Gamlin, 2006; Hattar et al., 2006; Klooster et al., 1995). Together with glutamate, ipRGCs co-utilise pituitary adenylate cyclase activating peptide (PACAP) acting via PAC1 receptors to excite its targets (Hannibal et al., 2017, 2000). It has been previously hypothesised that the OPN is involved in circadian timekeeping, due to its reciprocal connections with both the intergeniculate leaflet and the SCN (Gamlin, 2006; Klooster et al., 1995). It was also demonstrated that orexinergic fibres reach the OPN and display daily variation in orexin content (Chrobok et al., 2021b). However, daily changes in the spontaneous activity of OPN neurons or in their responsiveness to neuromodulators like orexin or PACAP has not been studied thus far.

Here, with the use of multi-electrode array (MEA) technology we report that OPN neurons increase their firing during the light phase. Additionally, we report for the first time the excitatory action of orexin A (OXA) upon the OPN neuronal activity, with the OX_2_ receptor mediating the effect. Interestingly, the responsiveness to OXA and the activation of PAC1 receptor by its selective agonist evoked stronger excitations during the night, when the spontaneous firing was lower. Altogether this report suggests intrinsic and extrinsic mechanisms to shape OPN neuronal activity in a daily fashion.

## 2. RESULTS

### 2.1. OPN neurons exhibit daily changes in spontaneous neuronal activity *ex vivo*

Neurons of the master clock and these localised in extra-SCN oscillators modulate their firing rates across the daily cycle to transmit information on their circadian phase (Hastings et al., 2018; Paul et al., 2020). Thus, we first aimed to establish, if OPN neurons similarly organise their neuronal activity in regards to the light-dark cycle. With the MEA technology *ex vivo*, we recorded spontaneous firing from nine pretectal slices, with four obtained from three rats culled at the middle of a day (ZT6), and five from three other rats culled at the opposite daily time point during the night (ZT18). Evaluation of neuronal activity from the recording locations in the OPN revealed higher firing rates during the day (4.17 ± 0.5 Hz, n=74), compared to the night (2.50 ± 0.3 Hz, n=102; *p*=0.0007, Mann-Whitney test; Fig. 1A,B). These results show that OPN neurons are more spontaneously active *ex vivo* during the light phase.

**Figure 1.**
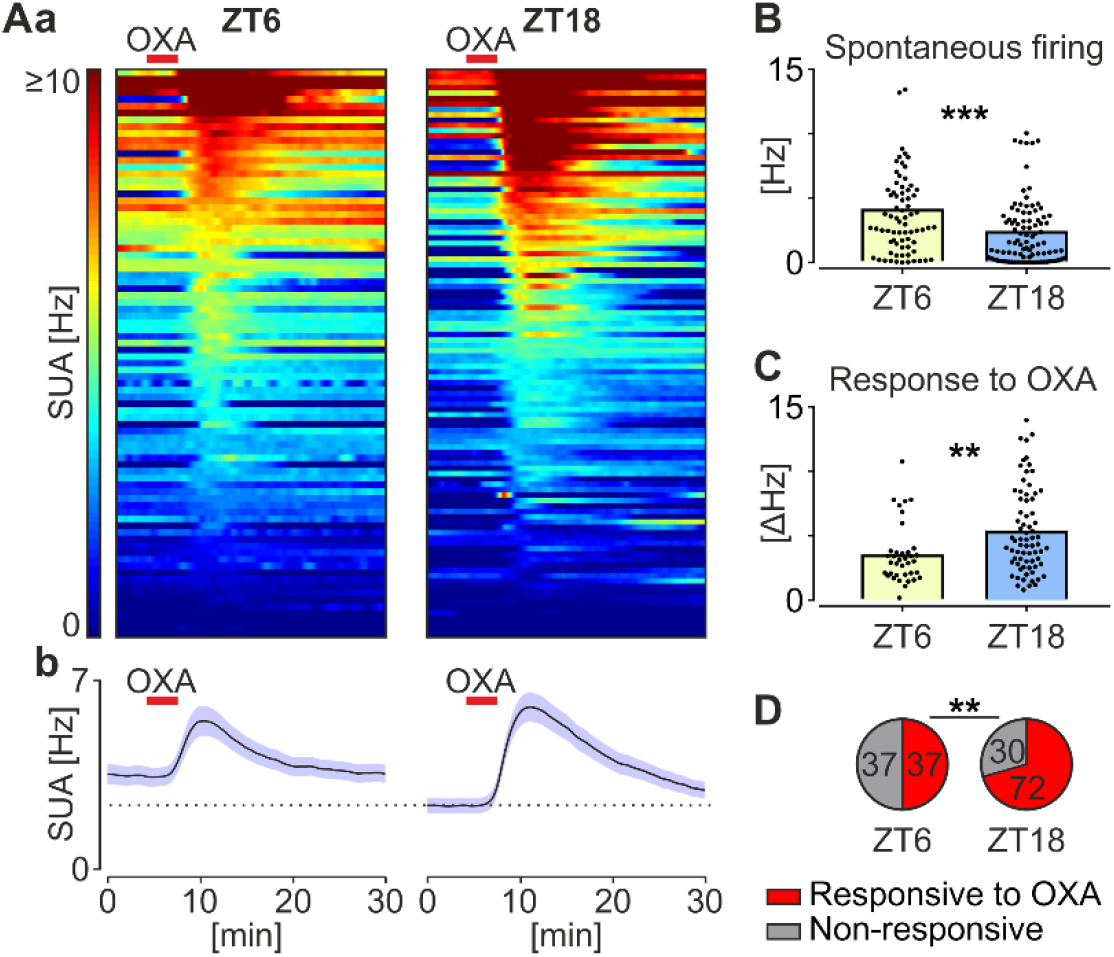
Neurons in the olivary pretectal nucleus (OPN) are more active during the day, but respond to orexin A (OXA) more robustly during the night. (**Aa**) Single unit activity (SUA) heatmaps showing the neuronal excitation following the application of OXA (200 nM) during the day (ZT6) and night (ZT18). Units were segregated from top to bottom, starting from these exhibiting the highest firing rates during the response to OXA. Warm colours code the highest, while cold – the lowest activity levels. Red bars represent the time of OXA application. (**Ab**) Average plots showing the mean ± SEM response to OXA at ZT6 and ZT18. Dotted line marks the baseline activity level at ZT18. (**B**) Spontaneous firing of individual OPN neurons. \*\*\**p*=0.0007, Mann-Whitney test. (**C**) The amplitude of individual responses to OXA. \*\**p*=0.0017, Mann-Whitney test. (**D**) Frequency of the response to OXA. \*\**p*=0.0074, Fisher’s test. ZT – Zeitgeber time.

### 2.2. OXA potently excites OPN neurons, with higher efficacy during the dark phase

The OPN has been shown to be a target of the orexinergic system, with orexinergic afferents showing a nocturnal rise in orexin content (Chrobok et al., 2021b). Hence, we next evaluated if orexinergic system may modulate neuronal activity in the OPN, by exogenous applications of OXA (200 nM) upon the same pretectal slices at day and night. Notably, OXA robustly activated OPN neurons, with higher amplitude seen during the night (ΔFR: 5.38 ± 3.3 Hz, n=37), compared to the day (ΔFR: 3.56 ± 2.3 Hz, n=72; *p*=0.0017, Mann-Whitney test; Fig 1A,C). Additionally, excitations were seen significantly more often at ZT18, compared to these at ZT6 (70.6% vs 50.0%, *p*=0.0074, Fisher’s test; Fig. 1D). This dataset suggests that the orexinergic system influences OPN neurons more effectively during the behaviourally active night.

### 2.3. OXA acts in the OPN predominately via OX_2_ receptor

Orexins bind to two metabotropic receptors: OX_1_ and OX_2_ receptor, which display a distinct expression pattern across the central nervous system (Li and de Lecea, 2020). To elucidate which of these two receptors mediates neuronal responses to orexins in the OPN, we applied OXA for the second time in the presence of a specific OX_2_ receptor antagonist – TCS-OX2-29 (10 µM) upon four pretectal slices previously treated with OXA in the standard ACSF. Then, OXA was re-applied for the third time, after the thorough washout of the antagonist. Two slices were recorded during the day and two during the night, and results were pulled together. Evidently, the presence of TCS-OX2-29 blocked the entirety of the response to OXA in the OPN (*p*<0.0001, n=49; Friedman’s test, Fig. 2A-D), with the response completely re-established after the antagonist washout (OXA1 vs OXA2+TCS: *p*<0.0001; OXA2+TCS vs OXA3: *p*<0.0001, Dunn’s multiple comparison test, Fig. 2D). This experiment shows the predominant contribution of OX_2_ receptor in the response of OPN neurons to OXA.

**Figure 2.**
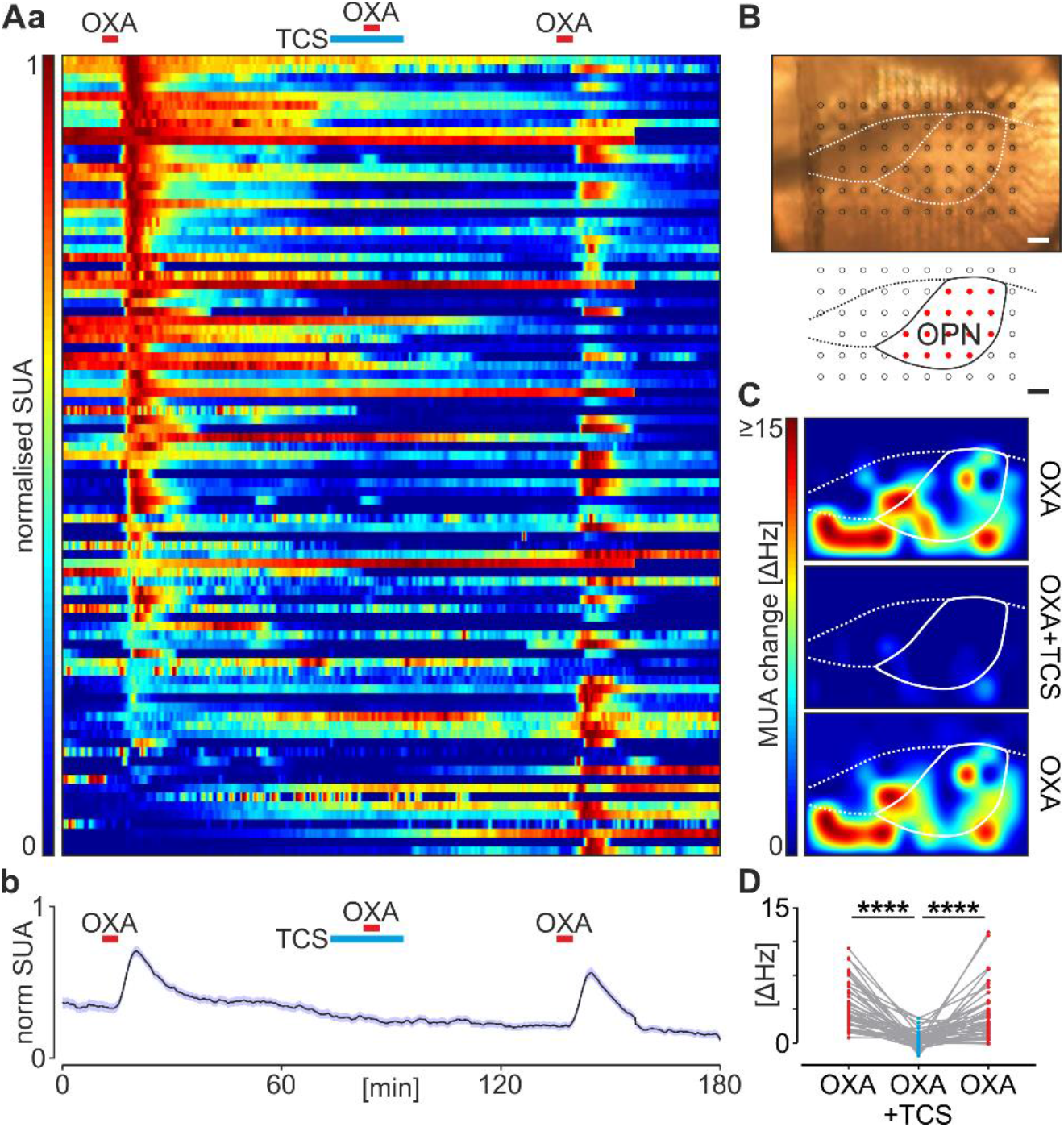
Orexin A (OXA) excites neurons in the olivary pretectal nucleus (OPN) via the OX_2_ receptor. (**Aa**) Single unit activity (SUA) heatmap normalised for each unit separately, showing the change in neuronal firing following the application of OXA (200 nM, red bar). Second application of OXA was performed in the presence of OX_2_ receptor antagonist – TCS-OX2-29 (TCS, blue bar). Finally, OXA was applied for the third time after the complete washout of the antagonist. Single units were segregated from top to bottom, with these showing the highest relative response to the first OXA application at the top. (**Ab**) Average plot of the mean ± SEM relative response to OXA in these three conditions. Note the lack of response to OXA in the presence of TCS. (**B**) Photomicrography showing the positioning of the OPN (outlined) on the multi-electrode array (grey circles), with the reconstruction below. Red circles code recording locations in the OPN. Bars show 100 µm. (**C**) Example spatial heatmaps of the multi-unit activity (MUA) showing the amplitude of the first, second (with TCS) and third (after TCS washout) response to OXA. (**D**) Changes in the response to OXA of individual OPN neurons in these three conditions. \*\*\*\**p*<0.0001, Dunn’s multiple comparison test.

### 2.4. Most of the OXA-sensitive neurons in the OPN are excited by the activation of PAC1 receptor

Retinal PACAP is released from the terminals of ipRGCs to excite neurons in the subcortical visual system via the PAC1 receptor (Hannibal et al., 2017, 2000). To test the possibility if orexinergic system targets the same OPN neurons that respond to the activation of PAC1 receptor, we applied maxadilan (MAX) – the specific PAC1 receptor agonist, following the application of OXA on five pretectal slices (two at ZT6 and three at ZT18). The application of MAX evoked an irreversible excitation of OPN neurons, with a higher amplitude during the night (ΔFR: 5.11 ± 0.5 Hz, n=21), compared to the day (ΔFR: 3.37 ± 0.4 Hz, n=26; *p*=0.0218, Mann-Whitney test; Fig 3A,B). The proportion of cells sensitive to the treatment was however similar between phases (*p*=0.5286, Fisher’s test; Fig 3C). Interestingly, 39.6% of neurons tested were activated by both neuropeptides, while 27.1% responded to OXA alone, and only 9.4% were exclusively excited by MAX (Fig 3D). These results suggest that OPN neurons sensitive to OXA also respond to retinal PACAP, with higher amplitudes of responses to PAC1 receptor activation observed during the dark phase.

**Figure 3.**
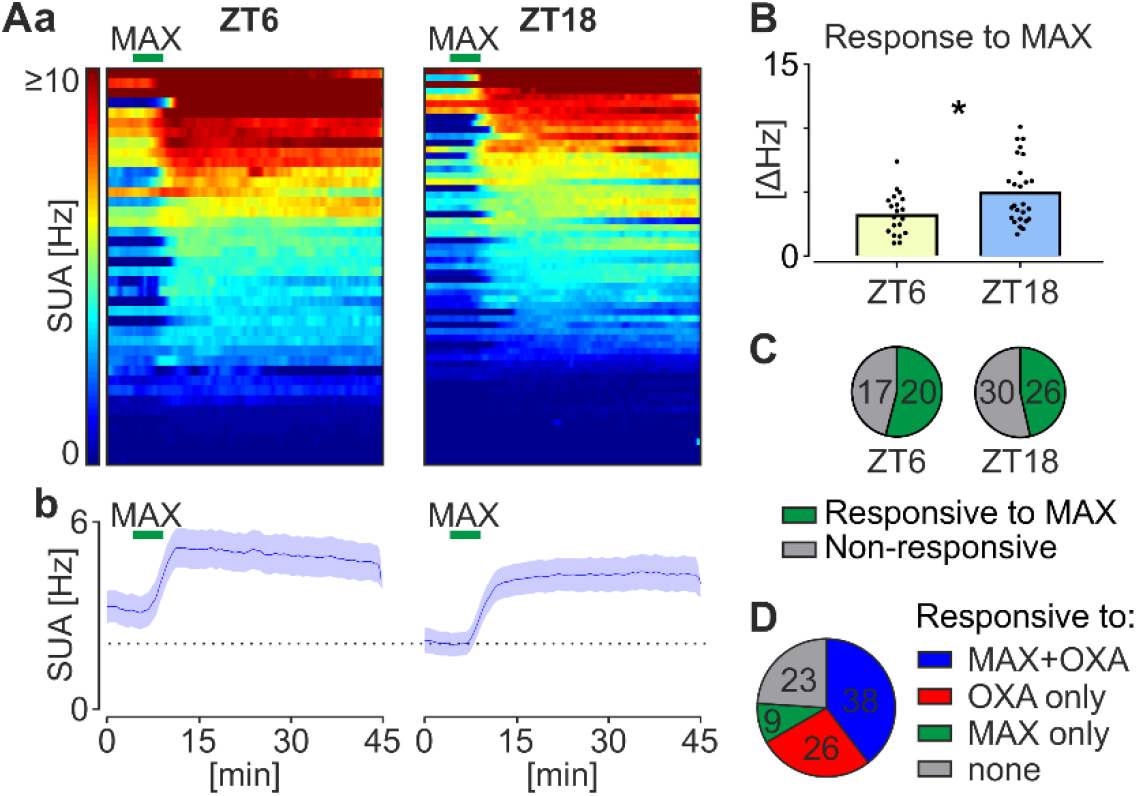
Most of the orexin A (OXA)-sensitive neurons in the olivary pretectal nucleus (OPN) are excited by the selective activation of PAC1 receptor with maxadilan (MAX). (**Aa**) Single unit activity (SUA) heatmaps showing the irreversible neuronal excitation following the application of MAX (100 nM) during the day (ZT6) and night (ZT18). Units were segregated from top to bottom, starting from these exhibiting the highest firing rates during the response to OXA. Green bars represent the time of MAX application. (**Ab**) Average plots showing the mean ± SEM response to MAX at ZT6 and ZT18. Dotted line marks the baseline activity level at ZT18. (**B**) The amplitude of individual responses to MAX. \**p*=0.0218, Mann-Whitney test. (**C**) Frequency of the response to MAX. ZT – Zeitgeber time. (**D**) The proportion of neurons responsive to MAX and OXA.

## 3. DISCUSSION

In this report we suggest that the orexinergic system of the lateral hypothalamus exerts excitatory actions upon OPN neurons predominately via OX_2_ receptor. Additionally, we identify day-time rise in the spontaneous firing rate of OPN neurons, oppositely phased to the enhanced sensitivity to OXA and the activation of PAC1 receptor, occurring at night. Finally, we propose a coupling of orexinergic and retinal information on the same OPN neurons, which respond to both OXA and PAC1 receptor agonist.

The modulation of neuronal activity in the subcortical visual system by orexins has been proposed before, for such areas as the lateral geniculate nucleus (LGN; all three parts) (Chrobok et al., 2021b, 2017, 2016; Orlowska-Feuer et al., 2019; Palus et al., 2015; Pekala et al., 2011), the SCN (Belle et al., 2014; Belle and Piggins, 2017; Brown et al., 2008), or recently for the superior colliculus (Chrobok et al., 2021a). The functional connection between these two systems has been also strengthened by reports on orexin action in the retina (Qiao et al., 2017; Zhang et al., 2018) and the primary visual cortex (Bayer, 2004). Here, we provide evidence for OXA to robustly excite the majority of OPN neurons. With the OPN being primarily responsible for the generation of a pupillary light reflex (Young and Lund, 1998), we suspect that orexins mediate direct effects of arousal on the pupil dilation. This hypothesis is reinforced by our observation, that OXA in the OPN acts through the OX_2_ receptor, suggested to transmit arousal-related actions of this peptide (Xu et al., 2004) and by the higher responsiveness of OPN neurons to OXA during the rats’ active phase.

Historically, the SCN was believed to exclusively generate circadian rhythmicity, at the level of clock gene expression and regulation of neuronal activity across the daily cycle (Takahashi, 2017). However, recent evidence strongly suggests that circadian control of many physiological processes must be devolved to local clocks localised in several brain structures and peripheral tissues (Abe et al., 2002; Begemann et al., 2020; Paul et al., 2020). Recently, we have reported circadian timekeeping properties also in the subcortical visual system outside of the SCN (Chrobok et al., 2021a, 2021b).

Here, we demonstrate that the OPN changes its firing rate between the light and dark phase, with higher activity levels recorded *ex vivo* during the behaviourally quiescent day; a time of a high retinal input *in vivo*. Interestingly, we furthermore reported higher nocturnal responsiveness to OXA and PAC1 receptor activation in the OPN. The first may be explained by improved sensitivity to orexins at night, when they are actually released *in vivo* due to an intensified behavioural activity and circadian regulation (Azeez et al., 2018). The higher night-time responsiveness to PAC1 receptor activation is however more surprising, as due to higher day-time activity of the retina, higher retinal PACAP release would be suspected during the light phase. This suggests an active mechanism promoting neuropeptidergic input from ipRGCs during the behaviourally active phase, when the retinal information is sparse but critically needed. To our knowledge, this is the first report on the daily variation of the OPN neurophysiology, but further detailed circadian studies are critically needed to resolve the intrinsic vs extrinsic source of this daily change.

## 4. EXPERIMENTAL PROCEDURE

### 4.1. Animals and ethical approval

The study described in this report was performed on six 6-7 week old male Sprague Dawley rats. Rats were bred and housed in the Animal Facility at the Institute of Zoology and Biomedical Research, Jagiellonian University in Krakow under standard 12:12 h light-dark cycle at 23 ± 2° C and 67 ± 3% relative humidity. All experiments were approved by Local Ethics Committee in Krakow and performed according to Polish regulations and the European Communities Council Directive (86/609/EEC). All possible efforts were made to minimise the number of animals used and their sufferings.

### 4.2 Electrophysiology

#### 4.2.1. Tissue preparation

Animals were anaesthetised with isoflurane (2 ml per anaesthetic chamber) and culled at two daily time points: in the middle of the day (ZT6), or in the middle of the night (ZT18). Then, 250 µm thick acute coronal pretectal slices containing the OPN were obtained in the same way as described previously for the LGN (Chrobok et al., 2021b). In brief, slices were cut on a vibroslicer (Leica VT1000S, Germany) in the ice-cold preparation artificial cerebro-spinal fluid (ACSF), composed of: NaHCO_3_ 25, KCl 3, Na_2_HPO_4_ 1.2, CaCl_2_ 2, MgCl_2_ 10, glucose 10, sucrose 125 and phenol red 0.01 mg/l. Then, slices were transferred to carbogenated recording ACSF (32° C, cooled down to room temperature), composed of (in mM): NaCl 125, NaHCO_3_ 25, KCl 3, Na_2_HPO_4_ 1.2, CaCl_2_ 2, MgCl_2_ 2, glucose 5, phenol red 0.01 mg/l. Sections were transferred to the recording chamber of the multi-electrode array (MEA) after a one hour incubation period.

#### 4.2.2. Recording

Neuronal activity of the OPN was evaluated with the use of the two-well perforated MEA *ex vivo* technology (Belle et al., 2021), according to a previously described procedure (Chrobok et al., 2021b, 2021c). Briefly, slices containing the OPN were transferred to the recording wells of the MEA2100-System (Multichannel Systems GmbH, Germany) and positioned upon the 6×10 recording array of the perforated MEA (60pMEA100/30iR-Ti, Multichannel Systems). Slices were constantly perfused with fresh recording ACSF (32° C, 2 ml/min) and sucked down the array. Data were continuously collected with Multi Channel Experimenter software (sampling frequency = 20 kHz; Multichannel Systems).

#### 4.2.3. Drugs

Orexin A (OXA, 200 nM; Bachem, Bubendorf, Switzerland), maxadilan (MAX; PAC1 receptor agonist; 100 nM; Bachem) and TCS-OX2-29 (TCS; OX_2_ receptor antagonist; 10 µM Tocris, Bristol, UK) were stored as 100 × concentrates at –20 ° C. All drugs were diluted in the fresh recording ACSF prior the application and delivered by bath perfusion. Approximately 4 min was needed for the drug to reach the recording chamber via the MEA tubing system.

#### 4.2.3. Spike-sorting and analysis

Data recorded with Multichannel Systems hardware and software were prepared for spike-sorting with the use of custom-made tools, described in detail in (Chrobok et al., 2021b, 2021c). Then, files were automatically spike-sorted with KiloSort programme (Pachitariu et al., 2016) in MatLab environment (R2018a version, MathWorks). Spike-sorting results were transferred into previously prepared CED-64 files (Spike2 8.11; Cambridge Electronic Design Ltd.) for further visualisation. Finally, spike-sorting results were manually revised in Spike2 8.11 with the aid of autocorrelation, principal component analysis and spike shape inspection to refine automatic sorting.

Further analyses were performed in Spike2 8.11. First, single unit activity (SUA) was 1 s binned. Then, mean spontaneous activity was explored in the 30 min epoch, starting 30 min after the initiation of recording. For examination of drug response, binned data were analysed in two epochs: 10 min before the drug application (baseline) and 30 min following the application (response). If after drug application the change in single unit activity (SUA) exceeded three standard deviations from baseline mean, a unit was classified as responsive. Response amplitudes were further calculated as a difference between 10 min baseline mean and the maximal value during the response to a drug.

#### 4.2.4 Data visualisation

Plots and pie charts were created in Prism 7 (GraphPad Software, USA). Heatmaps and average plots were generated in MatLab with custom-made scripts. For temporal heatmaps single unit data were first 10 s binned in NeuroExplorer 6 (Nex Technologies, USA) and then visualised as colour-coded firing rates or normalised SUA. Data normalisation was performed for each unit separately, with 1 coding the maximal and 0 – the minimal firing rate. Units were segregated from top to bottom with regards to the strength of the first response to a drug. Spatial heatmaps were generated from 100 s binned multi-unit activity (MUA) which was prepared in NeuroExplorer 6 for each recording location. The amplitude of response in MUA was calculated as the difference between a 100 s bin preceding the drug application and a 100 s bin during the maximal response.

### 4.3. Statistics

Statistical testing was performed in Prism 7. Differences between two unpaired groups were assessed with Mann-Whitney test, whereas Kruskal-Wallis test followed by Dunn’s multiple comparison test were used to test changes amongst three groups of paired data. Fisher’s test was used to study differences in group proportions.

